# H3K9me regulates heterochromatin silencing through incoherent feedforward loops

**DOI:** 10.1101/2023.11.02.565278

**Authors:** Kannosuke Yabe, Asuka Kamio, Satoyo Oya, Tetsuji Kakutani, Mami Hirayama, Yuriko Tanaka, Soichi Inagaki

## Abstract

Histone H3 lysine-9 methylation (H3K9me) is associated with condensed and transcriptionally inactive heterochromatin^1^. Although it has long been known that H3K9me silences transcription to control a wide variety of biological phenomena in many eukaryotic species^2,3^, how the silencing is regulated under the control of H3K9me is still largely unclear. Moreover, how cells delimit regions with H3K9me to avoid silencing essential genes remains unexplored. Here, using Arabidopsis genetic systems that induce H3K9me2 and its associated non-CG DNA methylation (mCH) in the transcribed region of genes *de novo*, we show that the accumulation of H3K9me2/mCH paradoxically also leads to the deposition of the euchromatic mark H3K36me3. This induction of H3K36me3 depends on a SET domain methyltransferase, ASHH3, and brings about anti-silencing by preventing the demethylation of H3K4me1 by LDL2, which mediates transcriptional silencing downstream of H3K9me2/mCH^4^. H3K9me2/mCH-driven antagonistic actions of ASHH3-H3K36me3 and LDL2-H3K4me1-loss also regulate the *de novo* silencing of reactivated transposable elements (TEs). These results demonstrate that H3K9me2 both facilitates and impedes silencing, and the incoherent feedforward loops fine-tune the fate of genes and TEs. Our results illuminate a novel elaborate mechanism for partitioning chromatin domains and provide insights into the molecular basis underlying natural epigenetic variation.

## Introduction

Ever since its discovery more than two decades ago, H3K9me has been regarded as the “heterochromatin mark”, which represses the transcriptional activity of underlying DNA^5^. H3K9me is associated with heterochromatin in many eukaryotes including metazoans, fungi, ciliates, and plants, and is involved in a variety of epigenetic phenomena, such as transposon silencing, imprinting, position effect variegation, and cell differentiation^2,3,6,7^. In some lineages of eukaryotes, such as vertebrates and plants, H3K9me is colocalized with DNA cytosine methylation (mC), which also plays a role in gene silencing^8,9^. In the plant *Arabidopsis thaliana*, H3K9 dimethylation (H3K9me2) and methylation at the CHG and CHH sites (H is A, T, or C; collectively referred to as mCH) are maintained through a self-reinforcing loop that is regulated by the H3K9 methyltransferases SUPPRESSOR OF VARIEGATION 3-9 HOMOLOGs (SUVH4/5/6) and DNA methyltransferases CHROMOMETHYLASEs (CMT2/3)^8^. Although the mechanisms to establish and maintain these heterochromatin marks in repressed regions of the genome have been elucidated^8,10^, how these marks silence transcription is largely unknown. In particular, the functions of these marks, aside from facilitating transcription silencing, have yet to be discovered.

Because heterochromatin formation is not a digital all-or-none phenomenon but rather a quantitative trait, different heterochromatin states have different dynamics and different genetic requirements^11–13^. Furthermore, the inherently spontaneous nature of heterochromatin formation necessitates mechanisms that continuously counteract heterochromatin^11,13,14^. Therefore, we focused on elucidating the mechanisms of establishing *de novo* heterochromatin formation and its counteraction using two experimental systems in which we investigated the early steps of heterochromatin (i.e., H3K9me2/mCH) formation in genes and TEs^15,16^. Our results revealed the unanticipated role of H3K9me2/mCH in counteracting heterochromatin silencing by recruitment of the euchromatic mark H3K36me3, which we propose is an intrinsic fail-safe mechanism that prevents the uncontrolled spreading of heterochromatin.

## Results

### Screening for factors that modulate transcription silencing

To elucidate the epigenome dynamics in heterochromatin silencing that is under the control of H3K9me2 and mCH in Arabidopsis, we leveraged a genetic mutant, *increase in BONSAI methylation 1* (*ibm1*), which ectopically accumulates H3K9me2/mCH in the transcribed regions (gene bodies) of more than three thousand active genes^15,17,18^. *ibm1*-induced H3K9me2/mCH in gene bodies causes a decrease in H3K4 monomethylation (H3K4me1) by the LSD1 family histone demethylase LDL2, which leads to transcriptional repression (Fig. 1a)^4^. Accordingly, the developmental phenotypes of the *ibm1* mutant are suppressed by the loss-of-function mutations of the *LDL2* gene due to the recovery of H3K4me1 and the gene expression. Here, we focused on the pathway by which H3K9me2/mCH induces a loss of H3K4me1 by LDL2 in gene bodies. We noticed that the extent of the decrease in H3K4me1 in the *ibm1* mutant showed great variation among different genes (Supplementary Fig. 1a) and thus reasoned that there may be additional factor(s) that modulate the LDL2-mediated silencing pathway (“Silencer” and/or “Anti-silencer”; Fig. 1a). To explore the underlying mechanisms, we searched for other chromatin features that are associated with the loss of H3K4me1 triggered by H3K9me2/mCH. With linear regression analyses of H3K9me2/mCH-induced H3K4me1 changes and each epigenomic feature, including H3K4me, H3K36me, H3K27me, H2B ubiquitination (H2Bub), H2A/H3 variants, and RNA polymerase II (RNAPII), we found that the genes with low levels of H3K36me3 and H2Bub showed tendencies to lose H3K4me1 in response to H3K9me2/mCH (Fig. 1b,c,d and Supplementary Fig. 1b,c). On the other hand, genes with higher levels of H2A.Z and H2A.W were more prone to the loss of H3K4me1 induced by H3K9me2/mCH (Fig. 1b and Supplementary Fig. 1b,c).

**Fig. 1:**
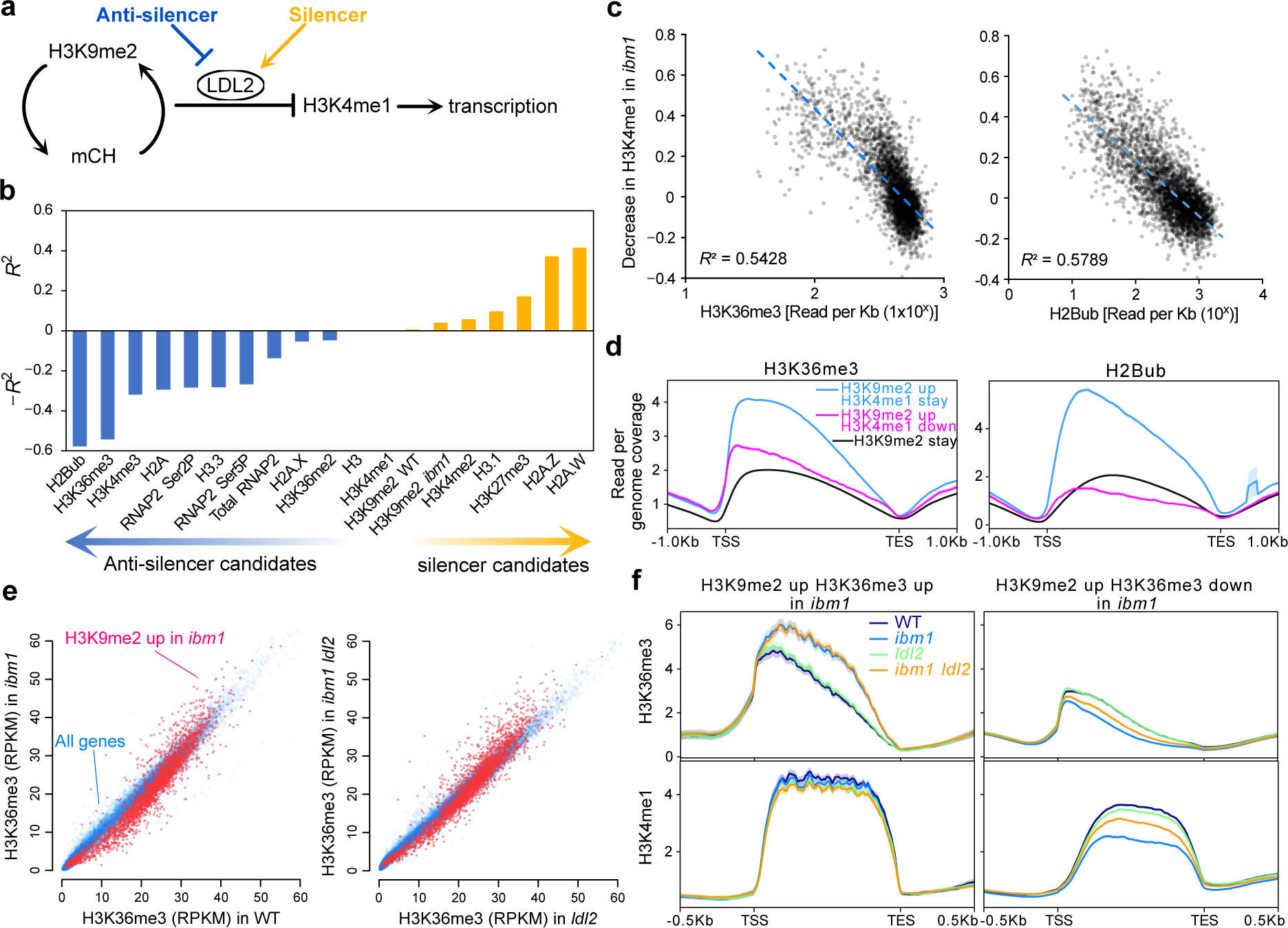
H3K36, as a putative anti-silencer, is induced by H3K9me2/mCH. **a**, Hypothetical factors that modulate silencing triggered by H3K9me2/mCH. **b**, Screening for other chromatin features that are correlated with the loss of H3K4me1 triggered by H3K9me2/mCH in *ibm1*. Pearson’s *R*^2^ values for factors positively correlated with the loss of H3K4me1 (silencer candidates) and -R^2^ values for factors negatively correlated with the loss of H3K4me1 (anti-silencer candidates) are shown. **c**, Correlations between H3K36me3 levels^20^ (left) and H2Bub levels^37^ (right), and the decrease in H3K4me1 in *ibm1* are shown as scatter plots, linear regression lines, and Pearson’s *R*^2^. Each dot represents a gene that accumulates H3K9me2 in *ibm1*. n = 3,395. **d**, Intragenic patterns of H3K36me3 and H2Bub in WT around genes categorized by H3K9me2 and H3K4me1 changes in *ibm1*. Among 3,449 genes that accumulate H3K9me2 in *ibm1* (“H3K9me2 up”), 743 genes showed clear decreases in H3K4me1 (“H3K4me1 down”), but others did not (“H3K4me1-stay”)^4^. **e**, H3K36me3 (Reads Per Kilobases per Million mapped reads (RPKM)) in *ibm1* and *ldl2* for all genes (blue) and genes accumulating H3K9me2 in *ibm1* (red). **f**, Intragenic patterns of H3K36me3 (top) and H3K4me1 (bottom) around genes categorized by H3K9me2 and H3K36me3 changes in *ibm1*.

### H3K9me2/mCH promotes H3K36me3 in gene bodies

From the above results, we hypothesized that H3K36me3 and/or H2Bub protect against the loss of H3K4me1 in *ibm1* (i.e., anti-silencer); thus, we first analyzed H3K36me3 and H2Bub patterns in *ibm1*, *ldl2*, and *ibm1 ldl2* double mutants by chromatin immunoprecipitation-sequencing (ChIP-seq). Many genes that gained H3K9me2 and lost H3K4me1 also lost H3K36me3 in *ibm1*, and this loss of H3K36me3 was partially suppressed by *ldl2* (Fig. 1e,f and Supplementary Fig. 2). This result is consistent with recent studies showing that H3K4me1 plays a role in recruiting the H3K36 methytransferase(s) to gene bodies^19,20^. Strikingly, however, a subset of genes that gained H3K9me2 also accumulated H3K36me3 (Fig. 1e,f), while H2Bub did not show this change (Supplementary Fig. 2). The accumulation of the euchromatic mark H3K36me3 triggered by H3K9me2 was rather unexpected and suggests a novel mechanism. The hyper-accumulation of H3K36me3 was not suppressed by *ldl2* (Fig. 1e,f), suggesting that H3K9me2 and/or mCH, but not the loss of H3K4me1, triggered the deposition of H3K36me3 (Fig. 2a). To rule out the possibility that IBM1 itself excludes H3K36me3 from genes independent of the function to remove H3K9me2/mCH, we analyzed the *ibm1 suvh4* double mutant that suppresses *ibm1*-induced H3K9me2/mCH^15,18^. The accumulation of H3K36me3 in *ibm1* was completely suppressed by *suvh4* (Supplementary Fig. 3), supporting the idea that H3K9me2/mCH paradoxically promotes the accumulation of H3K36me3 in gene bodies.

**Fig. 2:**
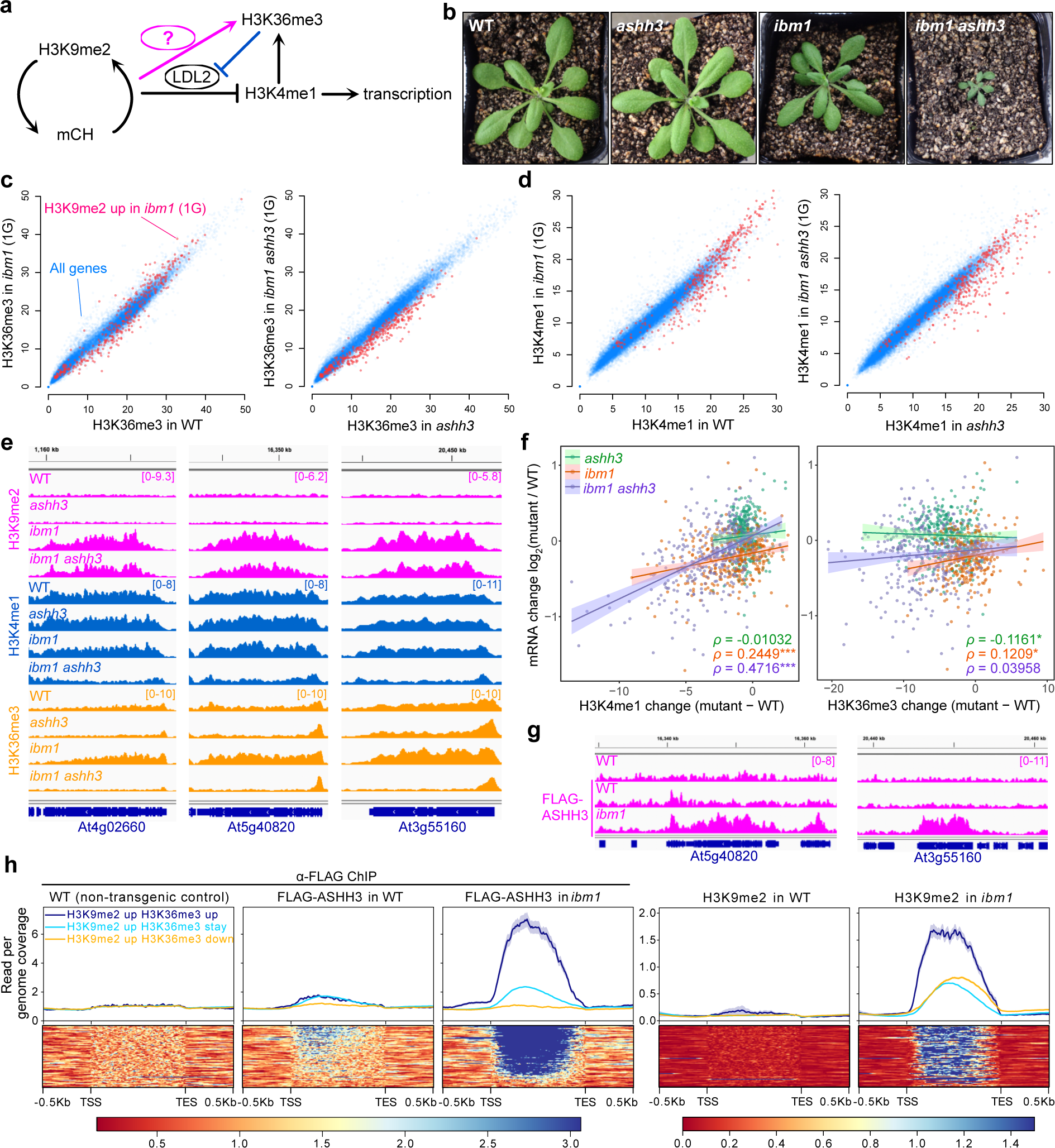
ASHH3 mediates H3K9me2/mCH-triggered H3K36me3 to prevents the loss of H3K4me1. **a**, A model depicting the H3K36me3 promotion by H3K9me2/mCH, which prevents the loss of H3K4me1 mediated by LDL2. **b**, Four-week-old WT, *ashh3*, *ibm1*, and *ibm1 ashh3* plants. **c**, H3K36me3 (RPKM) in *ibm1* and *ashh3* for all genes (blue) and genes with increased H3K9me2 in the 1st generation (1G) *ibm1* (red). **d**, H3K4me1 (RPKM) in *ibm1* and *ashh3* as in **c**. **e**, Browser views of H3K9me2, H3K4me1, and H3K36me3 in *ibm1* and *ashh3* around genes that have increased H3K9me2 and H3K36me3 in *ibm1*. **f**, Scatter plots comparing H3K4me1 changes (left) and H3K36me3 changes (right), with mRNA changes between WT and mutants. Spearman’s correlation coefficient p is shown for each WT-mutant comparison. *: 0.01 < p value < 0.05; ***: p value < 0.0001. **g**, FLAG-ASHH3 localization in WT and *ibm1* plants around genes with increased H3K9me2 and H3K36me3 in *ibm1* (shown in **e**). WT on top is the non-transgenic negative control. **h**, Average profiles (top) and heatmaps (bottom) of FLAG-ASHH3 and H3K9me2 around genes categorized by H3K9me2 and H3K36me3 changes in *ibm1*.

### ASHH3-mediated H3K36me3 protects against silencing

The genes that accumulated H3K36me3 in *ibm1* tended to retain H3K4me1 (Fig. 1f), suggesting again that H3K36me3 counteracts the loss of H3K4me1 mediated by LDL2 (Fig. 2a). To test this possibility and identify the H3K36 methyltransferase(s) responsible for H3K36me3 accumulation in *ibm1*, we made double mutants of *IBM1* and four predicted H3K36 methyltransferase genes, *ASHH1*, *2*, *3*, and *4*. Strikingly, we found that *ashh3* mutants strongly enhanced the developmental phenotypes of *ibm1*, while the other mutants did not (Fig. 2b and Supplementary Fig. 4), suggesting that ASHH3 is responsible for the induction of H3K36me3 in response to H3K9me2/mCH in *ibm1*. To test this hypothesis, we performed ChIP-seq for H3K4me1, H3K9me2, and H3K36me3 in *ibm1* and the *ibm1 ashh3* double mutant. The developmental phenotypes and sterility in *ibm1* progressively become severe across generations in response to the progressive accumulation of H3K9me2/mCH^18,21^, but *ibm1 ashh3* plants were already very defective in development and almost completely sterile in the 1st generation (Fig. 2b). Therefore, we performed ChIP-seq using the 1st generation of *ibm1* and *ibm1 ashh3*. In the 1st generation, ∼400 genes consistently accumulated H3K9me2 in both *ibm1* and *ibm1 ashh3* (Supplementary Fig. 5a). The increase in H3K36me3 was not caused by *ibm1* in the *ashh3* background (Fig. 2c), confirming that ASHH3 is responsible for H3K36me3 in response to H3K9me2/mCH. Furthermore, H3K4me1 was decreased in *ibm1 ashh3* compared to that in *ibm1* (Fig. 2d,e and Supplementary Fig. 5a,b), supporting the idea that ASHH3-mediated H3K36me3 protects against the loss of H3K4me1 that is mediated by LDL2. Interestingly, the loss of H3K4me1 in *ibm1 ashh3* (and in *ibm1* to a smaller extent) was significantly correlated with the downregulation of transcription, while the loss of H3K36me3 had a much weaker correlation (Fig. 2f). These results suggest that H3K4me1 is more critical for maintaining the transcriptional activity of genes than H3K36me3. Finally, we showed that genes that accumulate H3K9me2/mCH and H3K36me3 attract ASHH3 proteins in *ibm1* (Fig. 2g,h). Taken together, these results show that the accumulation of H3K9me2/mCH in gene bodies triggers ASHH3 recruitment and the consequent deposition of H3K36me3, which counteracts the loss of H3K4me1 mediated by LDL2 and transcriptional repression (Fig. 3a). The result that H3K36me3 was decreased in *ibm1 ashh3* compared to that in *ashh3* (Fig. 2c,e and Supplementary Fig. 5a,b) suggests that H3K4me1 also induces the deposition of H3K36me3, possibly by ASHH2, which recognizes H3K4me1 through the CW domain^19^.

**Fig. 3:**
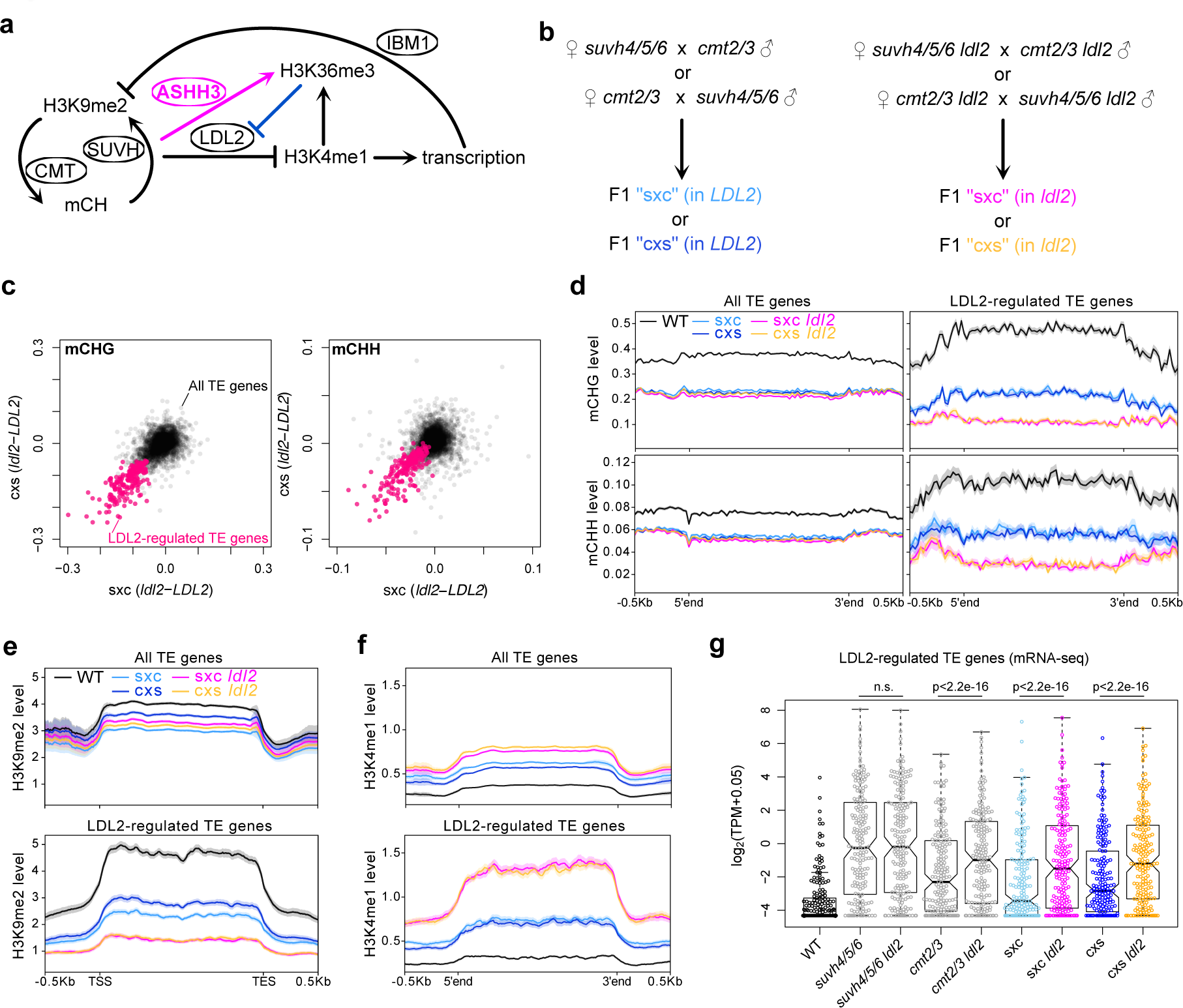
LDL2 facilitates TE silencing by removing H3K4me1. **a**, A model depicting the H3K9me2/mCH-promoted H3K36me3 by ASHH3, which prevents the loss of H3K4me1 mediated by LDL2. Transcribed sequences are targeted by IBM1 for H3K9 demethylation^15^. Therefore, the antagonistic actions of LDL2 and ASHH3 are predicted to affect H3K9me2/mCH dynamics through feedback regulation. **b**, Experimental design for analyzing the function of LDL2 in TE silencing. **c**, Effects of *ldl2* mutation on mCHG (left) and mCHH (right) establishment in sxc (x-axis) and cxs (y-axis). Each dot represents a TE-encoded gene (“TE gene”; n= 3,728), and red dots represent “LDL2-regulated TE genes” (n = 194), which show lower mCHG in sxc *ldl2* and cxs *ldl2* than in sxc *LDL2* and cxs *LDL2*, respectively (see methods). Averages of biological replicates are plotted. **d**, Averaged profiles of mCHG and mCHH in WT and four F1 lines around all TE genes (left) and LDL2-regulated TE genes (right). **e**, **f**, H3K9me2 (**e**), and H3K4me1 (**f**) patterns in WT and four F1 lines around all TE genes (top) and LDL2-regulated TE genes (bottom). **g**, mRNA levels (log_2_ of transcripts per million (TPM) + 0.05) of LDL2-regulated TE genes in WT, parental mutant plants, and F1 plants. Each circle represents each TE gene. In the boxplots, the centerline corresponds to the median, the notch represents the 95% confidence interval of the median, the upper and lower limits of the box correspond to the upper and lower quartiles, and the whiskers indicate the data range within 1.5x of the interquartile range (IQR). The *p* values are based on paired *t*-tests.

Because ASHH3 is not localized in TEs (Supplementary Fig. 6a), which have high H3K9me2/mCH levels, and ASHH3 accumulation in *ibm1* is positively correlated with H3K36me3 levels both in WT and *ibm1* (Supplementary Fig. 6b), we speculate that ASHH3 is recruited to the chromatin with both H3K9me2/mCH and H3K36me3, where it reinforces the amount of H3K36me3. This idea is consistent with results showing that genes that originally have higher levels of H3K36me3 tend to gain H3K36me3 (and escape from silencing) in *ibm1* (Fig. 1e,f). These results suggest that genes with high levels of H3K36me3 will be protected from silencing induced by H3K9me2/mCH (discussed later).

### Antagonistic LDL2-ASHH3 actions mediate the establishment of H3K9me2/mCH in TEs

The above results demonstrate that H3K9me2/mCH regulates transcription silencing through the combination of two incoherent feedforward loops (FFLs) (Fig. 3a and Supplementary Fig. 6c)^22^. Given that the downstream components in these FFLs (H3K36me3 and H3K4me1) are modulated by methylation and demethylation mechanisms from outside FFLs^19,20,23,24^, we speculate that incoherent FFLs, which are induced by H3K9me2/mCH, provide tunability to heterochromatin silencing and lead to different outcomes depending on the initial levels of H3K36me3 and H3K4me1; if loss of H3K4me1 occurs, the genes are silenced, and if the gain in H3K36me3 occurs, the genes remain transcribed. To elucidate whether this mechanism provides tunability to the silencing of TEs that are usually under the control of H3K9me2/mCH, we monitored the establishment of silencing by using genetic crosses between the triple mutant for the H3K9 methyltransferase genes *SUVH4*, *5*, *6* (*suvh4/5/6*) and the double mutant for the DNA methyltransferase genes *CMT2*, *3* (*cmt2/3*). Both *suvh4/5/6* and *cmt2/3* mutants lose H3K9me2 and mCH in TEs, but H3K9me2/mCH is largely recovered in their F1 hybrid, in which all the components of the H3K9me2-mCH feedback loop are present as heterozygous^16^. First, we conducted the genetic cross in backgrounds with or without the functional *LDL2* gene (Fig. 3b). In the *ldl2* background, the establishment of mCH in F1 was attenuated in many TEs compared with that in F1 in the *LDL2* background (Fig. 3c,d and Supplementary Fig. 7a-d), suggesting that LDL2 promotes mCH establishment. Accordingly, the H3K4me1 level was higher in these TEs in F1-*ldl2* than in F1-*LDL2*, and H3K9me2 showed the opposite trend (Fig. 3e,f and Supplementary Fig. 7e). These results suggest that LDL2 promotes H3K9me2/mCH establishment by removing H3K4me1 and repressing transcription. This hypothesis is consistent with our previous results that showed that IBM1 removes H3K9me2 from transcribed genes and TEs (Fig. 3a; ^15^). In agreement with this idea, the transcription levels of the TEs with attenuated H3K9me2/mCH in F1-*ldl2* were higher in F1-*ldl2* than in F1-*LDL2* (Fig. 3g).

We then tested the effects of ASHH3-H3K36me3, which counteracts LDL2, on mCH establishment in TEs by conducting the *suvh4/5/6*-*cmt2/3* cross in the *ashh3* background (Fig. 4a). We found several TEs that reproducibly had more mCH in the *ashh3* background than in the *ASHH3* background (Fig. 4b-d and Supplementary Fig. 8a-d and 9), suggesting that ASHH3 counteracts the accumulation of mCH as an anti-silencer in some TEs, as predicted by the model in Fig. 3a. Consistent with the elevated level of mCH in F1 in the *ashh3* background, H3K9me2 levels were also higher in these TEs in F1-*ashh3* than in F1-*ASHH3* (Fig. 4e and Supplementary Fig. 8e). These ASHH3-regulated TEs had higher H3K36me3 and H3K4me1 levels than other TEs in F1-*ASHH3*, and the levels of these marks were decreased in F1-*ashh3* (Fig. 4f,g and Supplementary Fig. 8e). Furthermore, we found that the transcription of most ASHH3-regulated TEs was significantly downregulated in F1-*ashh3* compared to F1-*ASHH3* (Fig. 4h and Supplementary Fig. 8f). Taken together, our results demonstrate that the antagonistic actions of LDL2 that removes H3K4me1 and ASHH3 that deposits H3K36me3, which are under the control of H3K9me2/mCH, mediates the *de novo* establishment of transcription silencing via intricate feedback and feedforward mechanisms (Fig. 3a).

**Fig. 4:**
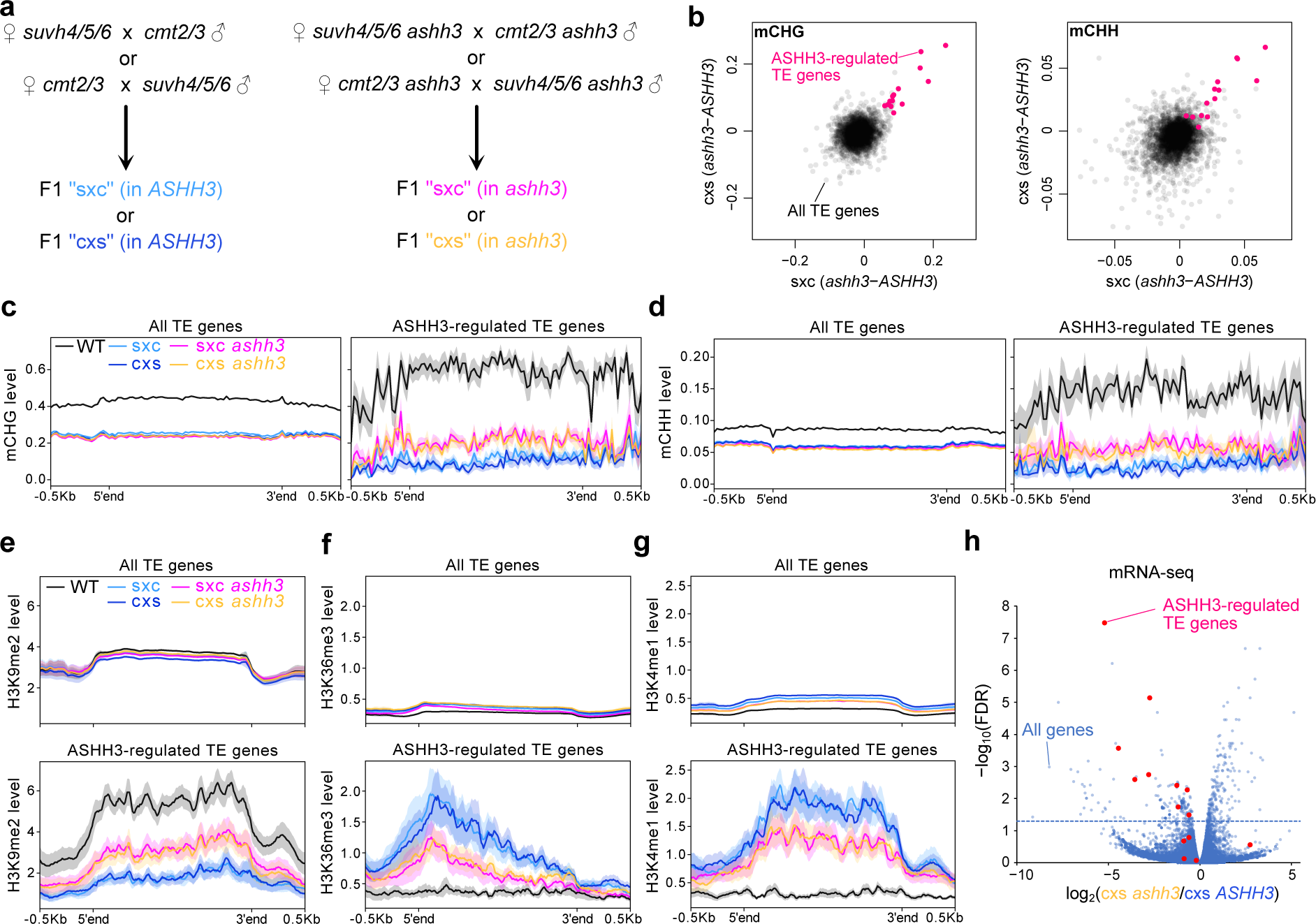
ASHH3 counteracts the silencing of several TEs. **a**, Experimental design for analyzing the function of ASHH3 in TE silencing. **b**, Effects of *ashh3* mutation on mCHG (left) and mCHH (right) establishment in sxc (x-axis) and cxs (y-axis). Each dot represents a TE gene (n = 3,551) and red dots represent “ASHH3-regulated TE genes” (n = 14), which show higher mCHG in sxc *ashh3* and cxs *ashh3* than in sxc *ASHH3* and cxs *ASHH3*, respectively (see methods). Averages of biological replicates are plotted. **c**,**d**, Averaged profiles of mCHG (**c**) and mCHH (**d**) in WT and four F1 lines around all TE genes (left) and ASHH3-regulated TE genes (right). **e**-**g**, H3K9me2 (**e**), H3K36me3 (**f**), and H3K4me1 (**g**) patterns in WT and four F1 lines around all TE genes (top) and ASHH3-regulated TE genes (bottom). **h**, Volcano plot of mRNA-seq comparing cxs *ashh3* and cxs *ASHH3*. Blue dotted lines represent FDR = 0.05. Blue dots represent all genes including protein-coding genes and TE genes (n = 33,603), and red dots represent ASHH3-regulated TE genes.

## Discussion

In this report, we demonstrated that H3K9me2/mCH both facilitates and impedes silencing, the latter of which is contrary to the widely believed canonical function of H3K9 methylation^3,5^. However, these two seemingly paradoxical functions of H3K9me2 can explain the robust differentiation of actively transcribed genes and silent TEs (Fig. 5). When H3K9me2/mCH accumulates in transcribed regions, which is thought to occur constantly but transiently^17,25^, genes with higher levels of H3K36me3 attract ASHH3 to methylate H3K36me3, which may repel LDL2 function and therefore protect H3K4me1 and transcription levels (Fig. 5a). H3K9me2/mCH is removed by transcription-promoted IBM1^15^. On the other hand, genes with less H3K36me3 cannot attract ASHH3 in response to H3K9me2/mCH accumulation; thus, LDL2 functions to remove H3K4me1, which eventually leads to transcriptional silencing and further accumulation of H3K9me2/mCH (Fig. 5b). We propose that this mechanism acts as a safeguard against the spurious silencing of essential genes. A recent report showed that certain genomic regions in human embryonic stem cells accumulate both H3K9me3 and H3K36me3^26^. Animals have multiple methyltransferases that catalyze H3K36me3, and the biological roles of some of these methyltransferases are still unknown^27^. Hence, it is tempting to speculate that the analogous mechanism(s) of fine-tuning heterochromatin silencing may also exist in animals.

**Fig. 5:**
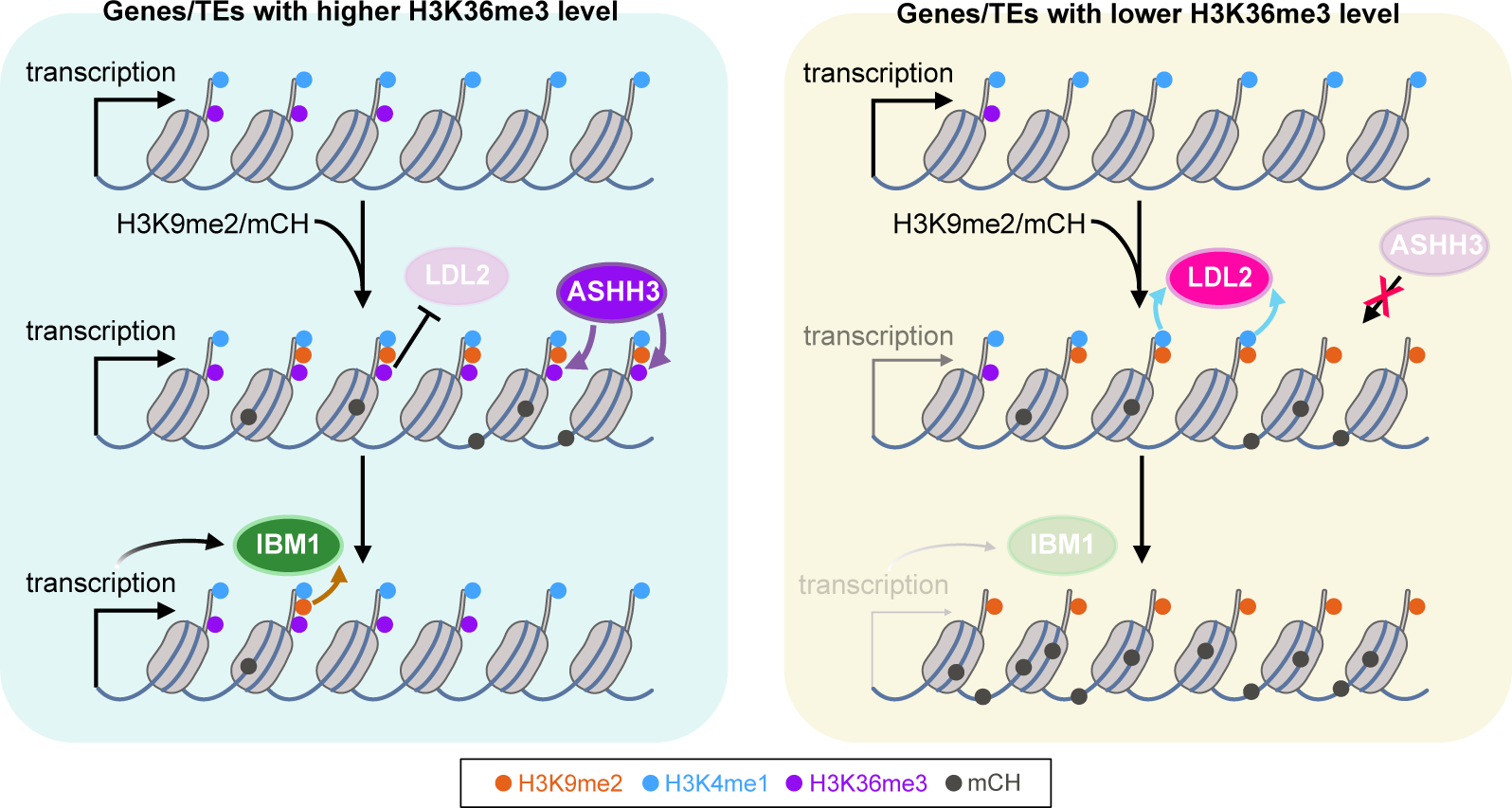
A model of the mechanism that partitions epigenetic marks by counteracting LDL2 and ASHH3. When H3K9me2/mCH accumulates in the transcribed region of genes or TEs, genes/TEs with high levels of H3K36me3 are kept in active chromatin status by the concerted functions of ASHH3 and IBM1 (left). Meanwhile, genes/TEs with lower levels of H3K36me3 cannot attract ASHH3, and thus silencing is established through the removal of H3K4me1 by LDL2 (right).

In animals and yeast, the similar noncanonical roles of H3K9 methylation in promoting gene transcription have been reported as mechanisms to solve the dilemma that transcription needs to occur for producing small-interfering RNA (siRNA) or PIWI-interacting RNA (piRNA), which are needed to maintain repressed heterochromatin status^28–32^. Plants have RNA polymerase IV dedicated to producing siRNA from repressed repeat sequences with H3K9me2/mCH, so that the above dilemma can be circumvented^33,34^. These mechanisms can be viewed as self-reinforcing feedback loops to stably maintain the repressed state of heterochromatin. Meanwhile, our study demonstrates a novel mechanism of incoherent FFLs, in which H3K9me2/mCH-promoted euchromatic mark H3K36me3 counteracts the establishment of silencing by inhibiting the demethylation of H3K4me1 (Fig. 5). Given that the deposition of H3K36me3 is coupled with the transcription process and H3K4me1, another transcription-promoted mark^19,20,23^, we anticipate that the H3K36me3-mediated anti-silencing mechanism uncovered here helps create various metastable epigenetic statuses depending on the underlying DNA sequence, transcription potential, and intracellular and extracellular environment; and thus, this mechanism could be a molecular basis for natural epigenetic variation^35,36^.

## Methods

### Plant materials and growth conditions

*Arabidopsis thaliana* strain Columbia-0 (Col-0) was used as the WT. *ibm1-4*^18^, *ldl2-1* (SALK_135831), *ldl2-2* (SALK_138820), *ashh3*/*sdg7-2* (SALK_131218C^38^), *ashh3*/*sdg7-4* (SALK_082736), *ashh3*/ *sdg7-5* (SALKseq_49152), *ashh3*/ *sdg7-6* (SALK_143603C), *ashh2*/*sdg8-1* (SALK_065480^39^), *ashh1*/*sdg26-1* (SALK_013895^40^), *ashh4*/*sdg24* (SK22803) and *suvh4/kyp* (SAIL588F05) are all in Col-0 background. The *ldl2-2* allele was used in the experiments in Fig. 1. The *ldl2-1* allele was used in the experiments in Fig. 3. For *ashh3*, the *ashh3/sdg7-2* allele was used in all epigenome analyses. Seeds were sown on Murashige and Skoog (MS) plates supplemented with 1% sucrose and solidified with gellan gum (Wako). Plants were grown in long-day conditions (16h light and 8h dark) at 22°C.

### Plasmid construction and transformation

pASHH3::3xFLAG-ASHH3-HA was constructed using NEBuilder HiFi DNA assembly Mix (New England Biolabs) and pPLV01 vector^41^. The promoter region of the *ASHH3* gene was amplified with the following primers: 5’- AATTCTAGTTGGAATGGGTTCTAAATCAATCATCAAACGATAAG-3’ and 5’- AATCTCCATCGTGATCTTTGTAATCCATGTCGAAAGAATCTAAAAG-3’. The coding region until just before the stop codon was amplified with the following primers: 5’- ACAAAGATGATGATGATAAGCCAGCCAGCAAAAAGGTC-3’ and 5’- TCTGGAACATCGTATGGGTAGACAATCTCCCAGTCTTCTC-3’ and then with the following primers: 5’- GATTACAAAGATCACGATGGAGATTACAAAGATCACGATATAGATTACAAAGATGATGAT GATAAG-3’ and 5’- GATCCTTATGGAGTTGGGTTTTAAGCGTAATCTGGAACATCGTATGGGTA-3’. The amplified fragments were assembled into pPLV01 using NEBuilder. The consequent plasmid was transformed into *Agrobacterium tumefaciens* GV3101::pMP90 containing pSOUP^42^ using electroporation. *ashh3* homozygous mutant plants were transformed with *Agrobacterium* with pASHH3::3xFLAG-ASHH3-HA using the floral dip method.

### Screening public dataset for silencing modifier

We used the following datasets. Histone H2A variants: H2A, H2A.W, H2A.X, and H2A.Z are from GSE50942^43^. H2B ubiquitination is from GSE112952^37^. H3 variants: H3.1 (HTR13) and H3.3 (HTR5) are from GSE34840^44^. H3K27me3, H3K36me2, and H3K36me3 are from PRJNA732996^20^. RNAPII (total), RNAPII-S5p, and RNAPII-S2p are from PRJDB10113^24^. Total H3, H3K4me1, H3K4me2, H3K4me3, and H3K9me2 in WT, and H3K9me2 in *ibm1* are from PRJDB5192^4^. All the sequence reads were mapped onto the TAIR10 genome using Bowtie^45^. The read count for each transcription unit was calculated using the “coverage” function of BEDTools^46^.

For correlation analysis, we used 3,395 protein-coding genes that accumulate H3K9me2 more in *ibm1* than in WT^4^. The Pearson correlation coefficient for each comparison between the explanatory variable (screening data set described above) and the decrease of H3K4me1 in *ibm1* (([H3K4me1]^WT^ - [H3K4me1]*^ibm^*^1^)/[H3K4me1]^WT^) as the objective variable was calculated. Metaplots of chromatin features around genes categorized by H3K9me2 and H3K4me1 changes in *ibm1* were drawn using the deeptools^47^.

### ChIP-seq

ChIP-seq for histone modifications was performed as described as eChIP-seq in ref^48^ with some modifications. ∼0.5 g of 2-week-old seedlings grown on MS plates were collected and frozen with liquid nitrogen, ground into a fine powder, and crosslinked with 1% formaldehyde in PBS buffer containing 1 mM Pefabloc SC (Merck), Complete proteinase inhibitor cocktail (Merck), and 0.3 % TritonX-100, for 10 min in room temperature. After quenching formaldehyde by adding 200 mM of glycine and incubating for 5 min, wash the fixed tissues with ice-cold PBS once with centrifugation at 5,000 g for 5 min. The tissue pellet was dissolved by the low-salt ChIP buffer without TritonX-100 (50 mM HEPES-KOH (pH 7.5), 150 mM NaCl, 1 mM EDTA, 0.1 % sodium deoxycholate, 0.1 % SDS) containing Complete proteinase inhibitor cocktail and sonicated with Picoruptor (Diagenode). After sonication debris was removed by centrifugation at 20,000 g for 10 min. The supernatant was transferred to a new tube and TritonX-100 was added to make the final concentration of 1 %. The sample was incubated with 1 μg of antibody: α-H3 (ab1791; Abcam), α-H3K9me2 (MABI0317; Wako), α-H3K4me1 (ab8895; Abcam), α-H3K36me3 (ab9050; Abcam), a-H2Bub (MM-0029; Medimabs). Antibody reaction was performed overnight at 4 °C with rotation. The sample containing the antibody-chromatin complex was incubated with Dynabeads Protein G (Thermo Fisher) for H3K9me2 or Dynabeads M-280 Sheep anti-Mouse IgG (Thermo Fisher) for other antibodies for 2h at 4°C with rotation. The beads were washed once with low-salt ChIP buffer (50 mM HEPES-KOH (pH 7.5), 150 mM NaCl, 1 mM EDTA, 1 % TritonX-100, 0.1 % sodium deoxycholate, 0.1 % SDS) containing Complete proteinase inhibitor cocktail, 2 times with high-salt ChIP buffer (50 mM HEPES-KOH (pH 7.5), 350 mM NaCl, 1 mM EDTA, 1 % TritonX-100, 0.1 % sodium deoxycholate, 0.1 % SDS), once with ChIP wash buffer (10 mM Tris-HCl (pH 8.0), 250 mM LiCl, 0.5 % NP-40, 1 mM EDTA, 0.1 % sodium deoxycholate), and once with TE buffer. The chromatin was eluted by adding ChIP elution buffer (50 mM Tris-HCl (pH 7.5), 10 mM EDTA, and 1 % SDS) and incubating at 65 °C for 15 min. Subsequently, 4 μl of Proteinase K (20 mg/ml; Thermo Fisher) was added to the sample and incubated at 55 °C overnight. The immunoprecipitated DNA was purified using the Monarch PCR & DNA Cleanup Kit (New England Biolabs). The libraries for Illumina sequencing were constructed using the ThruPLEX DNA-Seq Kit (Clontech) and purified using SPRIselect beads (Beckman Coulter). The sequencing was performed by the HiSeqX or NovaSeqX Plus sequencer (Illumina).

ChIP-seq for 3xFLAG-ASHH3 was performed with some modifications from histone modifications eChIP-seq. 0.8 g of seedlings were collected and crosslinked in 1 % formaldehyde solution by applying vacuum for 15 min. Glycine was added to the solution and further incubated for 5 min with vacuum. Tissues were washed with water, dried with paper towels, frozen with liquid nitrogen, and kept in a -80 °C freezer until use. The fixed tissue was ground into fine powder and proceeded as described above. The sonication was performed with Covaris S220 Focused-ultrasonicator (Covaris). Immunoprecipitation was conducted with 1pg of a-FLAG antibody (F3165; Merck).

Reads were mapped on the Arabidopsis TAIR10 genome using Bowtie^45^. The read count for each transcription unit was calculated using the “coverage” function of BEDTools^46^. Metaplots and heatmaps were drawn using the deeptools^47^.

### mRNA-seq

Total RNA was extracted from 2-week-old seedlings grown on an MS plate using RNeasy Plant Mini Kit (Qiagen) following the manufacturer’s instructions. mRNA-seq libraries were constructed from 600ng of total RNA using the KAPA mRNA HyperPrep Kit (Kapa Biosystems). Three independent biological replicates were analyzed for each genotype. The resulting libraries were 150 bp paired-end sequenced by the HiSeqX or NovaSeqX Plus sequencer (Illumina).

Reads were mapped on the Arabidopsis TAIR10 genome using STAR aligner^49^ with --outFilterType, BySJout; -- alignSJoverhangMin, 8; -- alignSJDBoverhangMin, 1; -- clip3pNbases, 50; -- quantMode GeneCounts parameters. The resulting per-gene counts were used for downstream analysis. For TE gene analysis (Fig 3g, 4h), the strand that showed larger counts was chosen for each TE gene based on the sum of all samples and used for further analyses because some TE genes show higher transcription levels of antisense strand than sense strand.

### Enzymatic Methyl-seq

Whole genome DNA methylation sequencing was performed using the NEBNext Enzymatic Methyl-seq (EM-seq) Kit (New England Biolabs) following the manufacturer’s instructions. Genome DNA was extracted from 2-week-old seedlings grown on an MS plate using Nucleon PhytoPure (Cytiva). Approximately 400 ng of DNA was fragmented by sonication using a Covaris S220 Focused-ultrasonicator (Covaris) and size-selected to enrich 400-600 bp fragments using SPRIselect beads (Beckman Courter). EM-seq libraries were constructed from 30ng of fragmented DNA. Two independent biological replicates were analyzed for each genotype. The resulting libraries were 150 bp paired-end sequenced by the HiSeqX sequencer (Illumina).

Reads were trimmed for the adapter sequences and low-quality regions using the Trimmomatic program^50^. Subsequent trimmed reads were mapped to the *Arabidopsis* reference genome TAIR10 using Bismark ver. 0.10.1^51^ with -n 1 -l 20 -e 90 parameters. Deduplication and methylation extraction were also performed using Bismark. Counting all the methylated unmethylated C in each transcription unit was conducted using the “map” function of BEDTools^46^. The methylation level was calculated as the ratio of total methylated cytosines over total cytosines in each region (weighted methylation level^52^). The LDL2-regulated TE genes (Fig. 3c) were extracted with the following criteria. (1) Having more than 100 counts for total CHG sites in both biological replicates of all samples of cxs *LDL2*, cxs *ldl2*, sxc *LDL2*, and sxc *ldl2*. (2) ([mCHG] in cxs *ldl2*) - ([mCHG] in cxs *LDL2*) and ([mCHG] in sxc *ldl2*) - ([mCHG] in sxc *LDL2*) in both biological replicates (in total 4 comparisons) are all below -0.05. The ASHH3-regulated TE genes (Fig. 4b) were extracted with the following criteria. (1) Having more than 100 counts for total CHG sites in both biological replicates of all samples of cxs *ASHH3*, cxs *ashh3*, sxc *ASHH3*, and sxc *ashh3*. (2) ([mCHG] in cxs *ashh3*) - ([mCHG] in cxs *ASHH3*) and ([mCHG] in sxc *ashh3*) - ([mCHG] in sxc *ASHH3*) in both biological replicates (in total 4 comparisons) are all more than 0.05. Metaplots were drawn using the deeptools^47^.

## Data availability

The high-throughput sequencing data generated in this study is available in the NCBI Gene Expression Omnibus (GEO) database under the accession number GSE246527.

## Acknowledgements

We thank Mayumi Takahashi, Hideko Watabe, and Juliarni for technical assistance and Arabidopsis Biological Resource Center for mutant seeds. This study was supported by grants from Japan Science and Technology Agency (JST) PRESTO (no. JPMJPR17Q1 to S.I.), Japan Society for the Promotion of Science (JSPS) (nos. JP20H05913 and JP22H02299 to S.I. and JP21H04977 to T.K.), and Mitsubishi Foundation (no. 202110005 to S.I.).

## Author contributions

S.I. conceived and designed the study. K.Y., A.K., S.O., and T.K. contributed to the design of the experiments. K.Y., A.K., S.O., M.H., Y.T., and S.I. conducted the experiments. K.Y., A.K., and S.I. analyzed the data. S.I. wrote the manuscript incorporating other author’s comments.

## Competing interests

The authors declare no competing interests.

Correspondence and requests for materials should be addressed to Soichi Inagaki.

## Legends for Supplementary Figures

**Supplementary Fig. 1: Screening for silencing modifier(s). a**, Relationships between changes in H3K9me2 and H3K4me1 levels (RPKM) within each gene in *ibm1* compared to WT. Data is from ref 4. **b**, Correlations between chromatin features (x-axis), and the decrease of H3K4me1 in *ibm1* (y-axis) are shown as scatter plots, linear regression lines, and Pearson’s *R*^2^. Each dot represents each gene that accumulates H3K9me2 in *ibm1*. n = 3,395. **c**, Intragenic patterns of chromatin features around genes categorized by H3K9me2 and H3K4me1 changes in *ibm1*. Among 3,449 genes that accumulate H3K9me2 in *ibm1* (“H3K9me2 up”), 743 genes showed clear decreases of H3K4me1 (“H3K4me1 down”) but others did not (“H3K4me1-stay”)^4^.

**Supplementary Fig. 2: Histone modification patterns in *ibm1* and *ibm1 ldl2*. a**, Scatter plots of H3K9me2, H3K4me1, H2Bub, and H3 in *ibm1* and *ldl2* for all genes (blue) and genes accumulating H3K9me2 in *ibm1* (red). **b**, Intragenic patterns of H3K9me2, H2Bub, and H3 around genes categorized by H3K9me2 and H3K36me3 changes in *ibm1*.

**Supplementary Fig. 3: H3K36me3 pattern in *ibm1 suvh4*. a**, Intragenic patterns of H3K36me3 (top) and H3 (bottom) around genes categorized by H3K9me2 and H3K36me3 changes in *ibm1* and TE genes. **b**, Browser views around genes with increased H3K36me3 in *ibm1*, which was suppressed in *ibm1 suvh4*.

**Supplementary Fig. 4: Effect of *ashh* mutations on *ibm1* phenotype.** Four-week-old plants with combinatorial mutations of *ibm1* and *ashh1/2/3/4*. The gene names in capital letters and lower letters indicate wild-type allele and loss of function allele, respectively. *ashh3-2* allele is the same allele as Fig. 2b but the different individuals. *ashh3-4/5/6* are other alleles with independent T-DNA insertion events (see methods for material information).

**Supplementary Fig. 5: Histone modification patterns in *ibm1* and *ibm1 ashh3*. a**, Intragenic patterns of H3K36me3, H3K4me1, H3K9me2, and H3 in *ibm1* (1G) and *ashh3* around genes categorized by H3K9me2 and H3K36me3 changes in *ibm1*, and TE genes. **b**, Heatmaps of H3K36me3, H3K4me1, H3K9me2, and H3 in *ibm1* (1G) and *ashh3* around 74 genes that are increased in H3K9me2 and H3K36me3 levels in *ibm1* (1G).

**Supplementary Fig. 6: ASHH3 localization and incoherent feedforward loops. a**, Intragenic patterns of FLAG-ASHH3 around genes categorized by H3K9me2 and H3K36me3 changes in *ibm1*, and TE genes. WT is used as the non-transgenic negative control. Two independent FLAG-ASHH3 transgenic lines were analyzed both in WT and *ibm1* background. **b**, The relationship between H3K36me3 level in WT (left) and *ibm1* (right), and ASHH3 hyper-accumulation in *ibm1*. **c**, Four types of incoherent FFLs^22^ in the box and two incoherent FFLs revealed in this study (right).

**Supplementary Fig. 7: LDL2-regulated establishment of TE silencing. a**, Effects of *ldl2* mutation on mCG in sxc (x-axis) and cxs (y-axis). Each dot represents each TE gene (n= 3,728) and red dots represent “LDL2-regulated TE genes” (n = 194), which show lower mCHG in sxc *ldl2* and cxs *ldl2* than in sxc *LDL2* and cxs *LDL2*, respectively (Fig. 3c). Averages of biological replicates are plotted. **b**, Averaged profiles of mCG in WT and four F1 lines around all TE genes (left) and LDL2-regulated TE genes (right). **c**,**d**, mCG (left), mCHG (middle), and mCHH (right) levels (ratios of methylated to all cytosines in each context) of LDL2-regulated TE genes (**c**) and all TE genes (**d**) in WT, parental mutant plants, and F1 plants. Each circle represents each TE gene. In the boxplots, the center line corresponds to the median, the notch represents the 95% confidence interval of the median, the upper and lower limits of the box correspond to the upper and lower quartiles, and the whiskers indicate the data range within 1.5x of the IQR. The *p*-values are based on paired *t*-tests. **e**, H3K9me2 (left), H3K4me1 (middle), and H3 (right) levels of LDL2-regulated TE genes in WT and F1 plants. The *p* values are based on paired *t*-tests.

**Supplementary Fig. 8: Effect of *ashh3* on the establishment of TE silencing. a**, Effects of *ashh3* mutation on mCG in sxc (x-axis) and cxs (y-axis). Each dot represents each TE gene (n= 3,551) and red dots represent “ASHH3-regulated TE genes” (n = 14), which show higher mCHG in sxc *ashh3* and cxs *ashh3* than in sxc *ASHH3* and cxs *ASHH3*, respectively (Fig. 4b). Averages of biological replicates are plotted. **b**, Averaged profiles of mCG in WT and four F1 lines around all TE genes (left) and ASHH3-regulated TE genes (right). **c**,**d**, mCG (left), mCHG (middle), and mCHH (right) levels (ratios of methylated to all cytosines in each context) of ASHH3-regulated TE genes (**c**) and all TE genes (**d**) in WT, parental mutant plants, and F1 plants. In **c**, each circle represents each TE gene, and the horizontal bars indicate the medians. The *p*-values are based on paired *t*-tests. In the boxplots (**d**), the center line corresponds to the median, the notch represents the 95% confidence interval of the median, the upper and lower limits of the box correspond to the upper and lower quartiles, and the whiskers indicate the data range within 1.5x of the IQR. (**e**), H3K9me2, H3K4me1, H3K36me3, and H3 levels of ASHH3-regulated TE genes in WT and F1 plants. The *p*-values are based on paired *t*-tests. **f**, Volcano plots of mRNA-seq comparing WT and cxs *ASHH3* (left), and WT and cxs *ashh3* (right). Blue dotted lines represent FDR = 0.05. Blue dots represent all genes including protein-coding genes and TE genes (n = 33,603), and red dots represent ASHH3-regulated TE genes.

**Supplementary Fig. 9: TEs anti-silenced by ASHH3.** Browser views showing DNA methylation in CG, CHG, CHH contexts, H3K9me2, H3K36me3, H3K4me1, and H3 around two ASHH3-regulated TE genes, At2g45230 (top) and At5g47815 (bottom).

