## Supplementary Figures for "H3K9me regulates heterochromatin silencing through incoherent feedforward loops"

### Supplementary Fig. 1

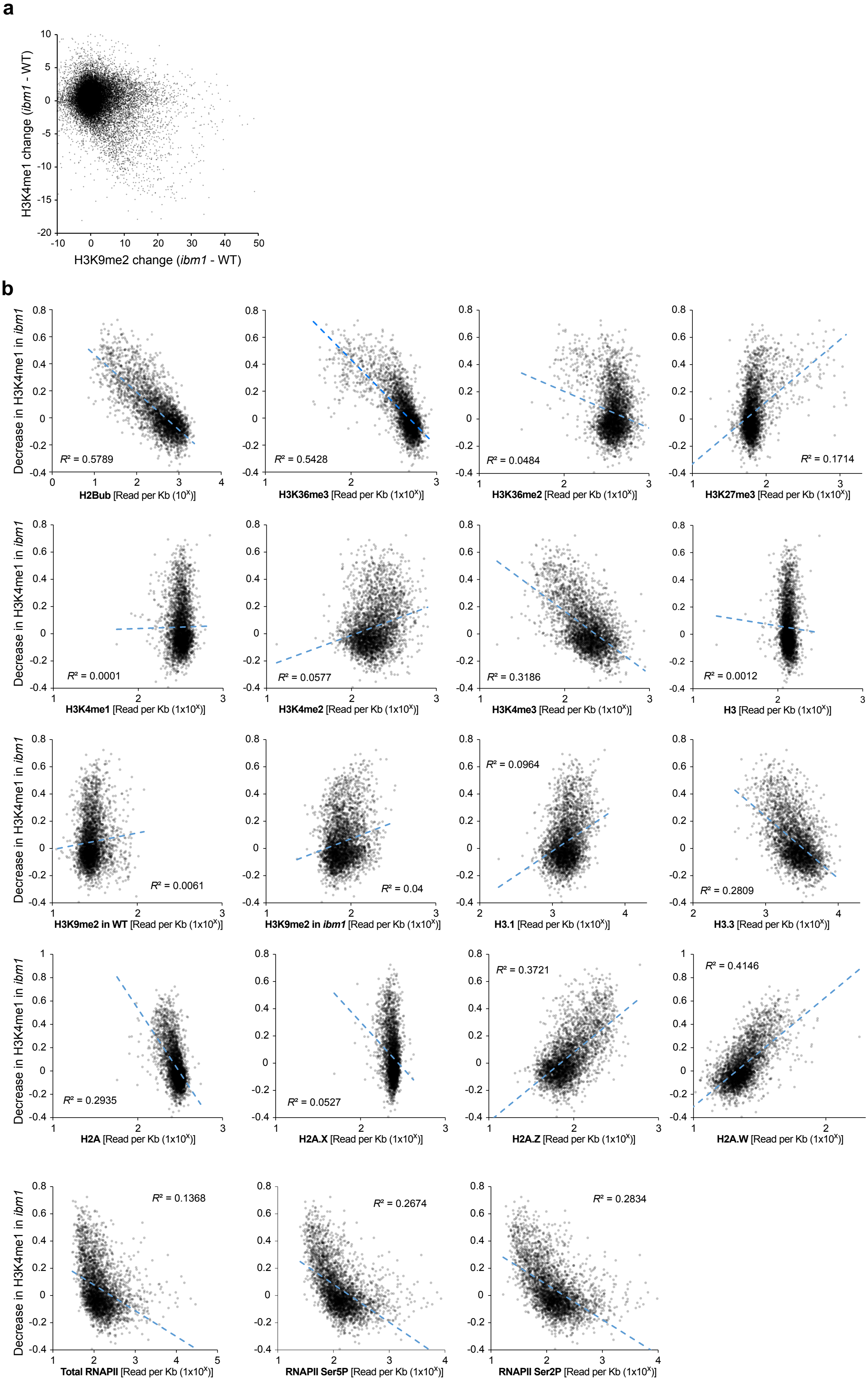

Supplementary Fig. 1 (continued)

c

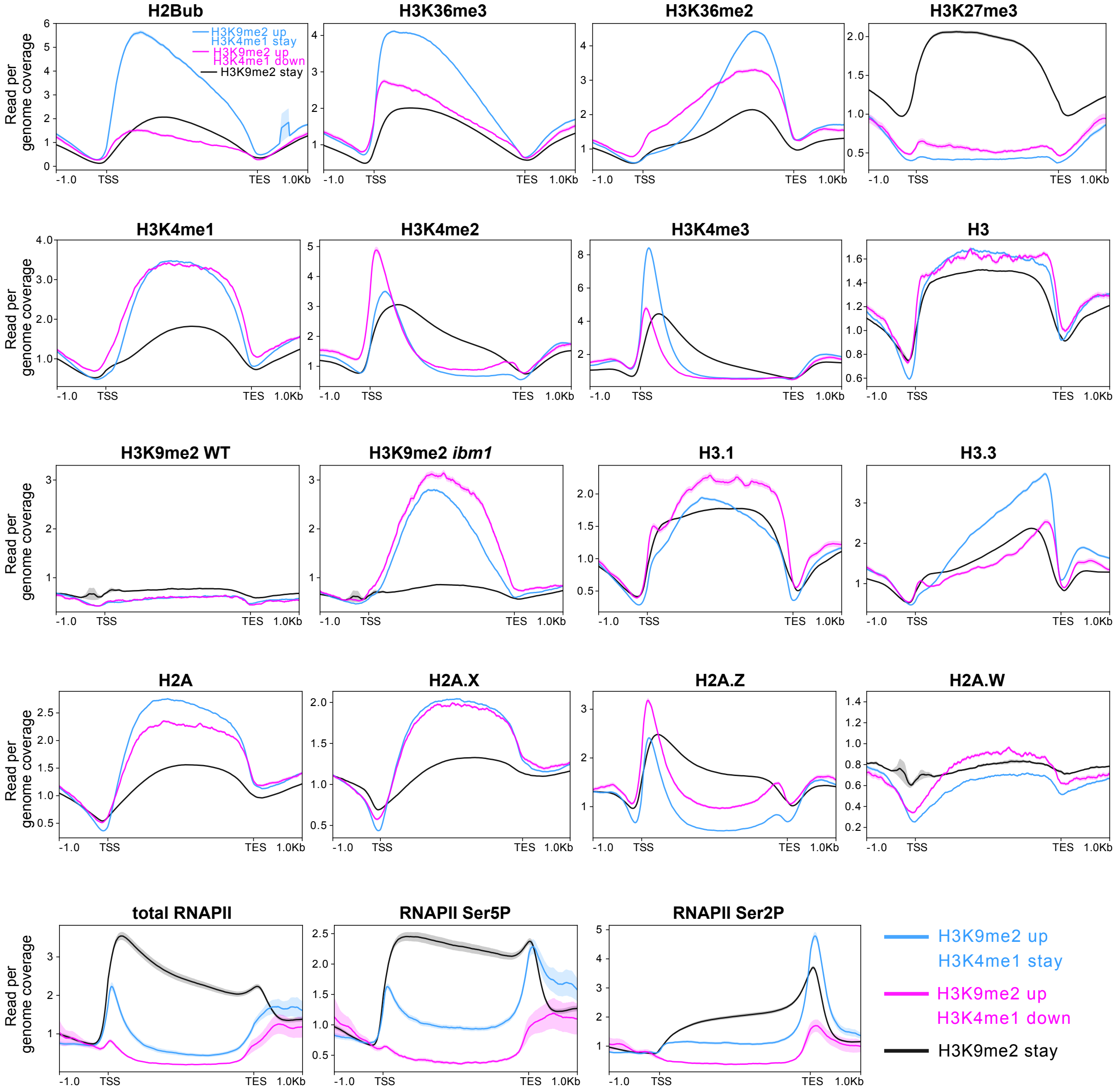

#### Supplementary Fig. 2

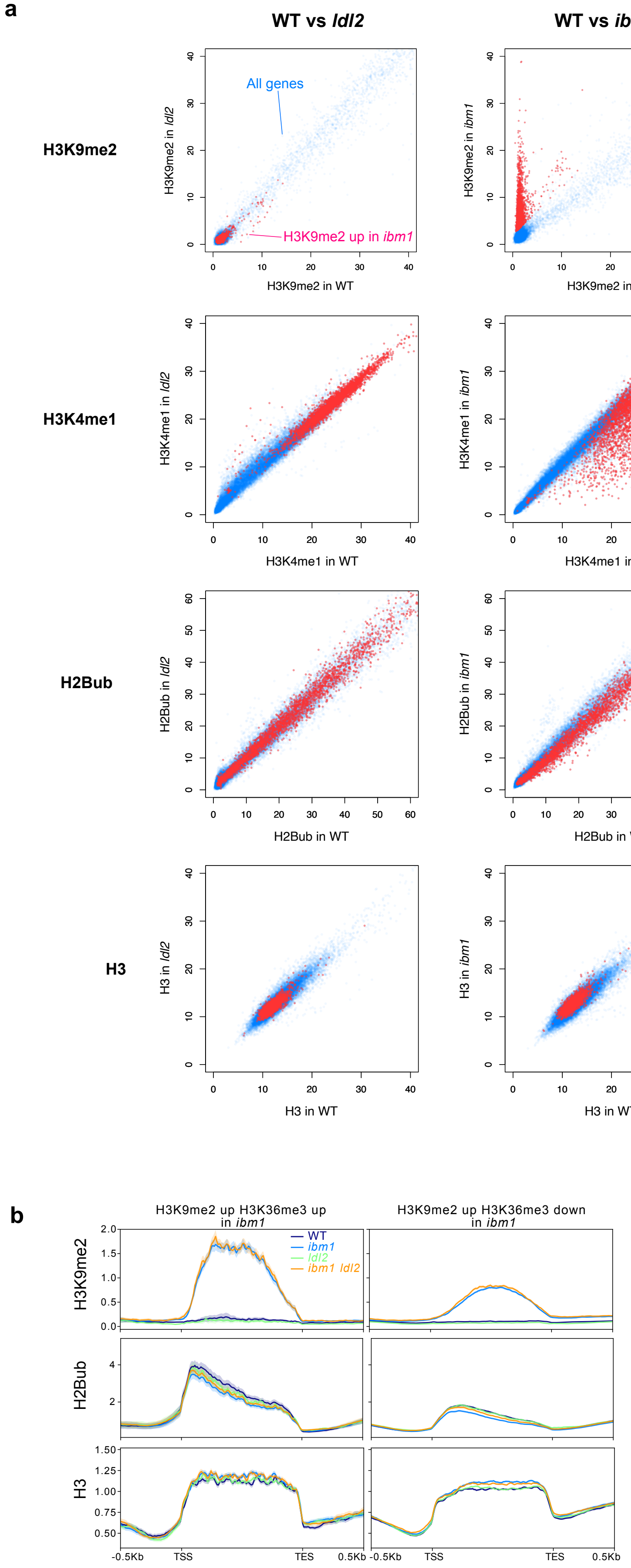

#### Supplementary Fig. 3

**a**

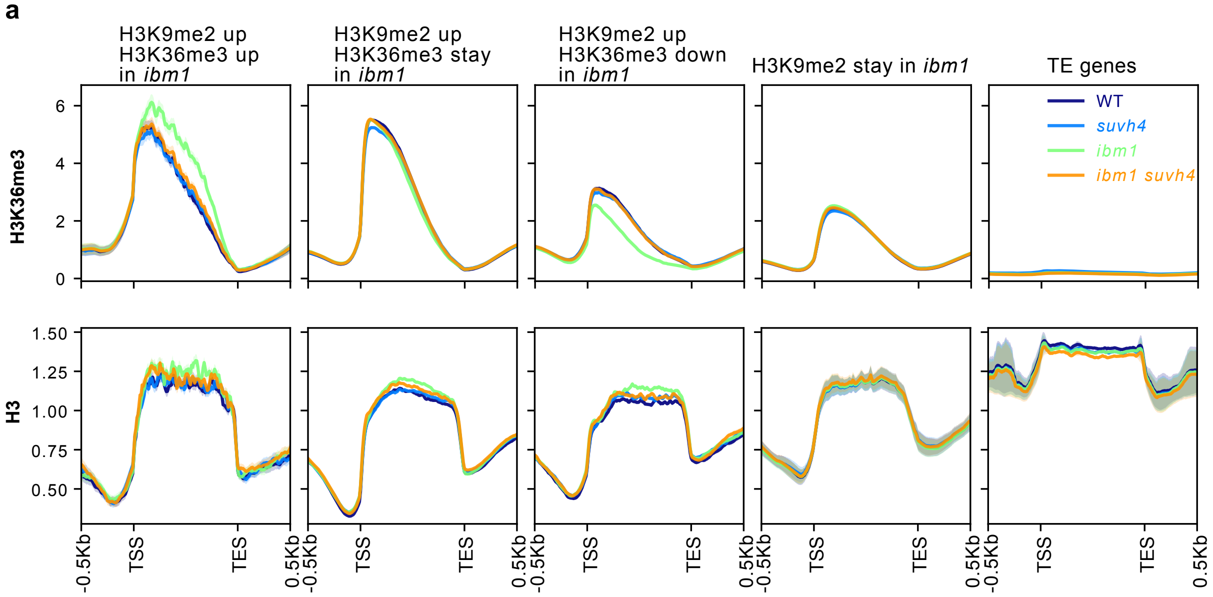


**b**

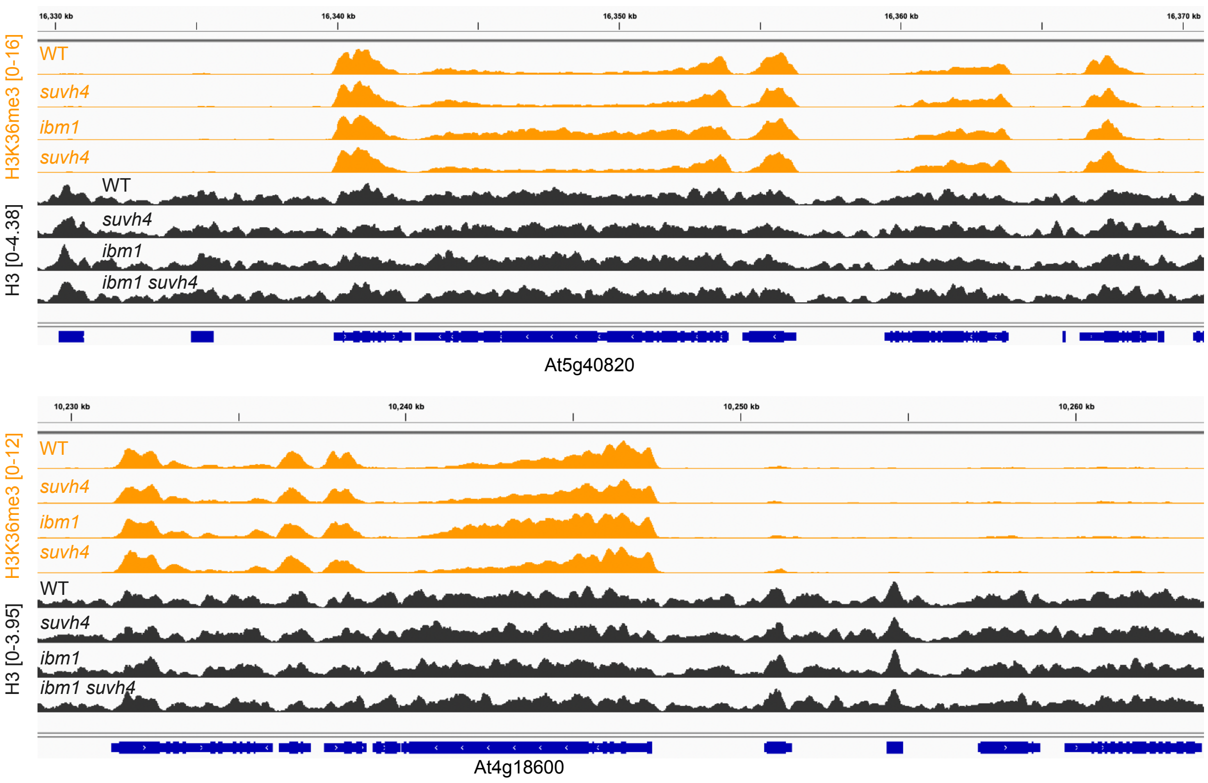

**Supplementary Fig. 4**

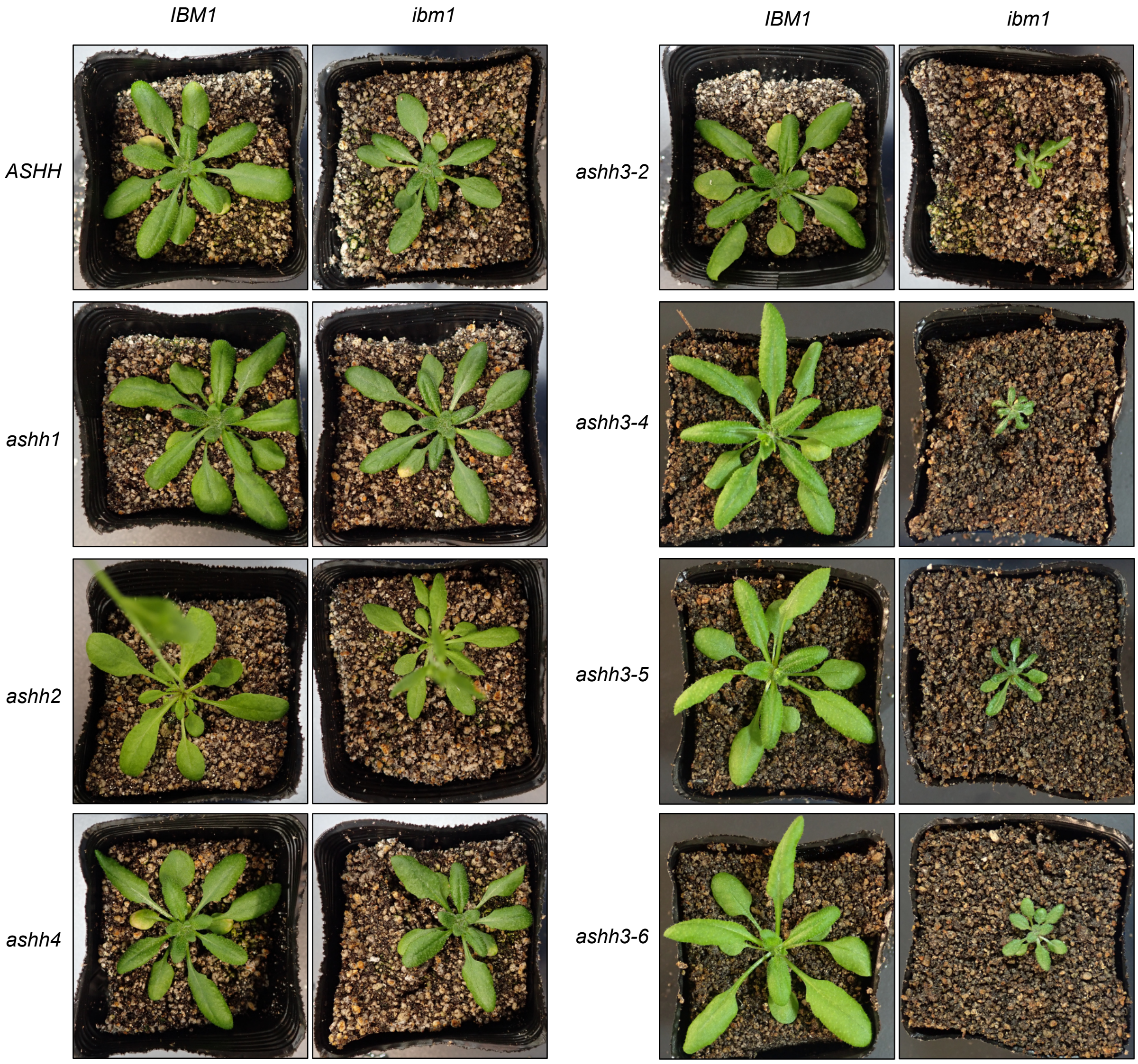

#### Supplementary Fig. 5

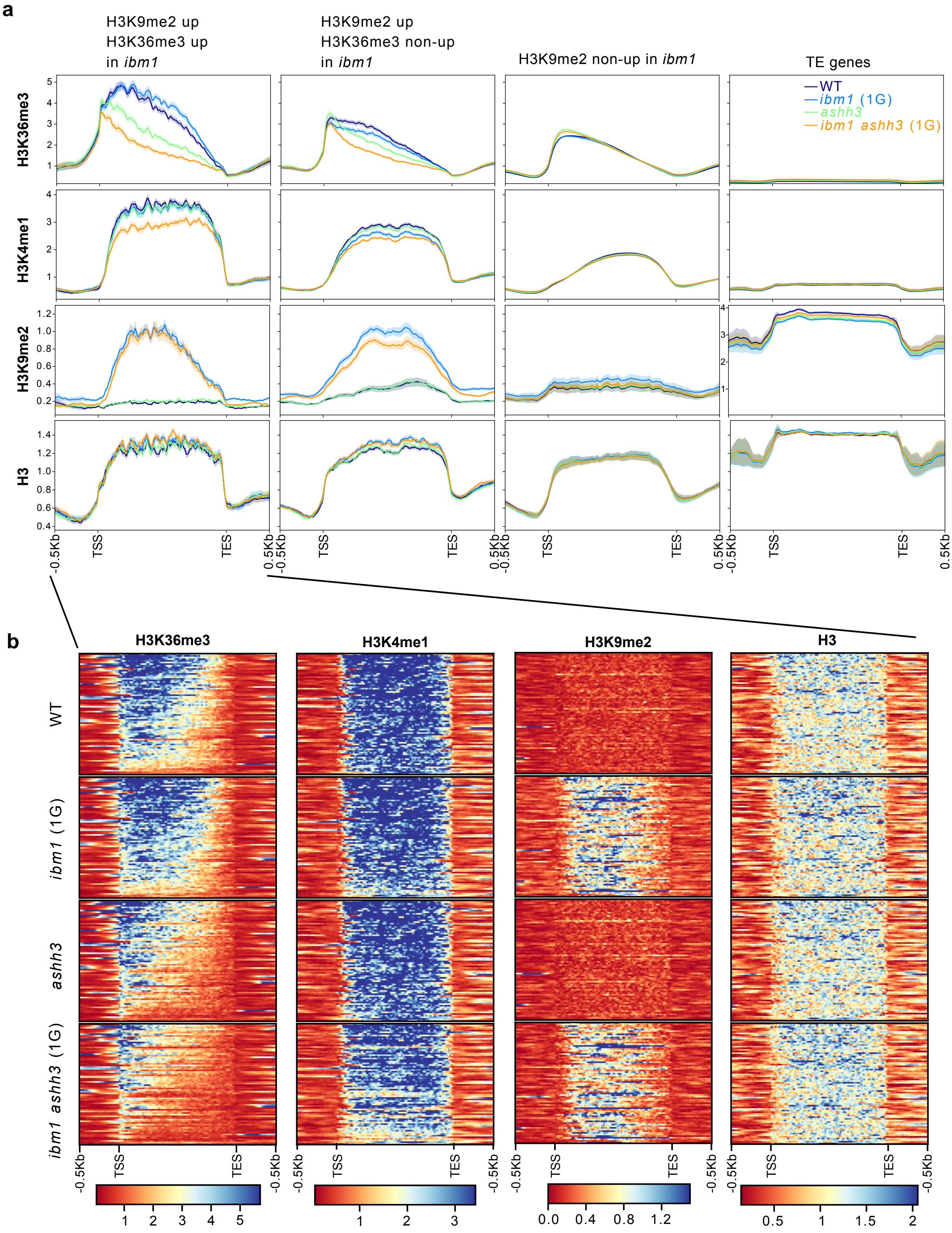

**Supplementary Fig. 6**

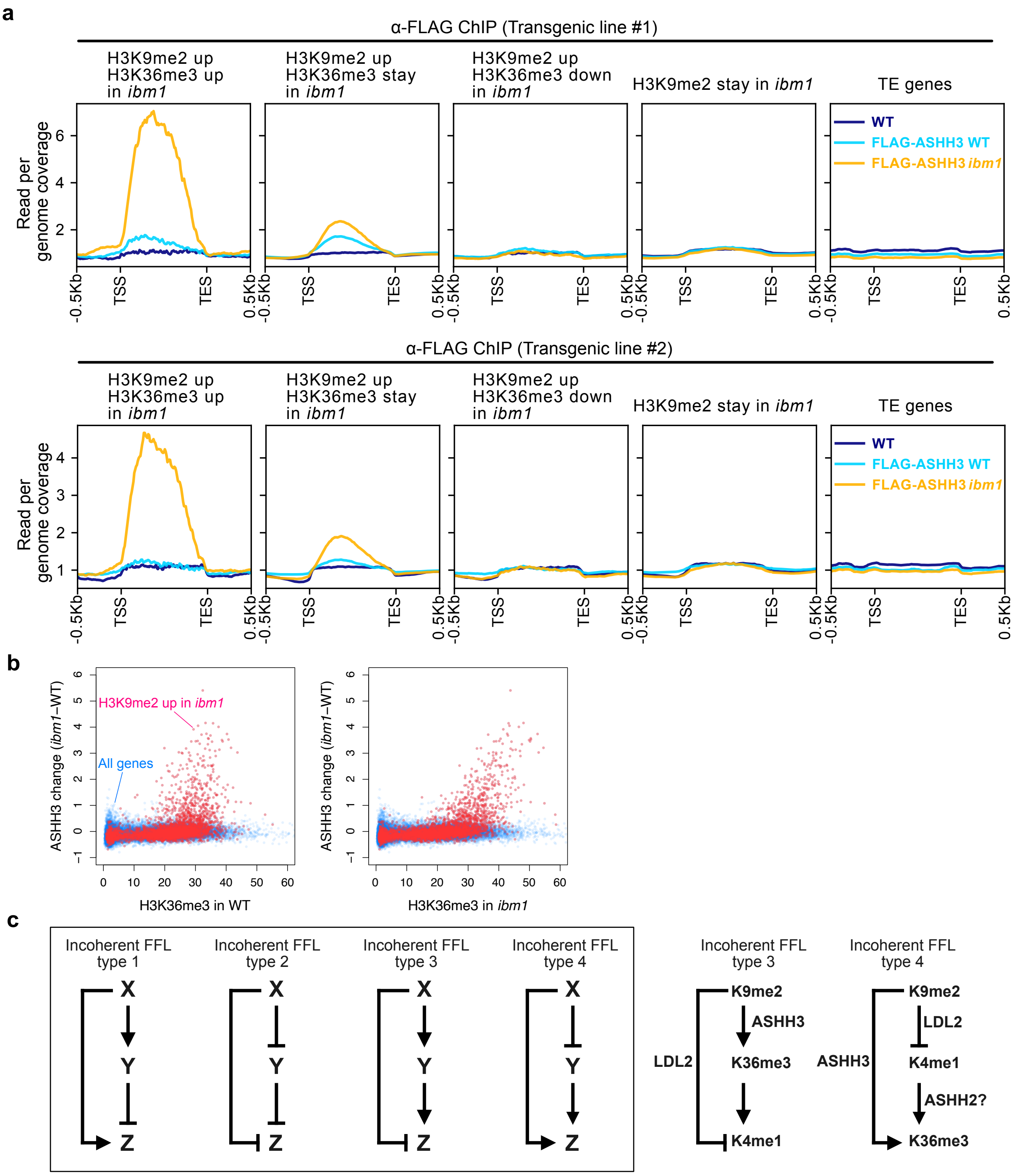

#### Supplementary Fig. 7

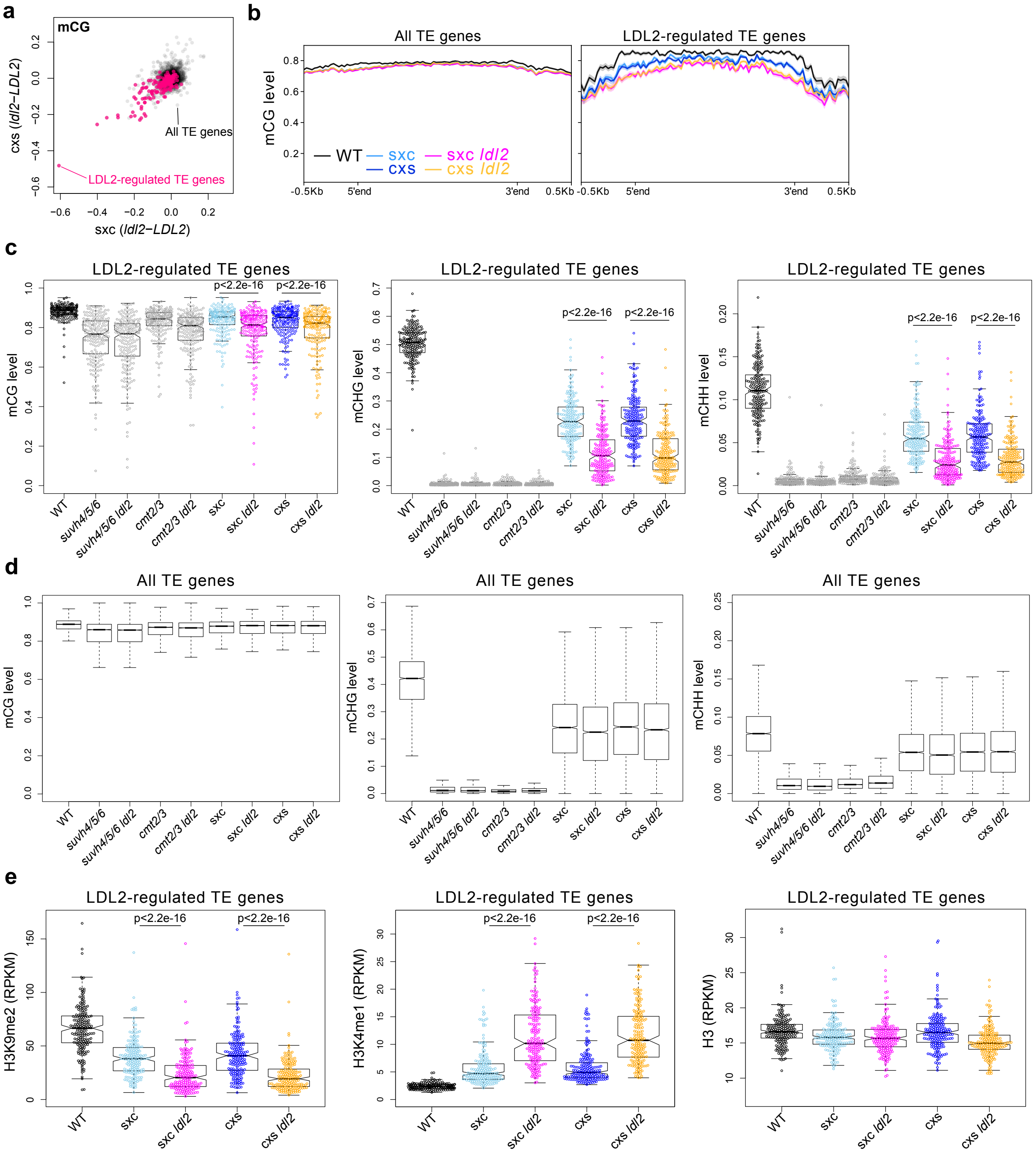

#### Supplementary Fig. 8

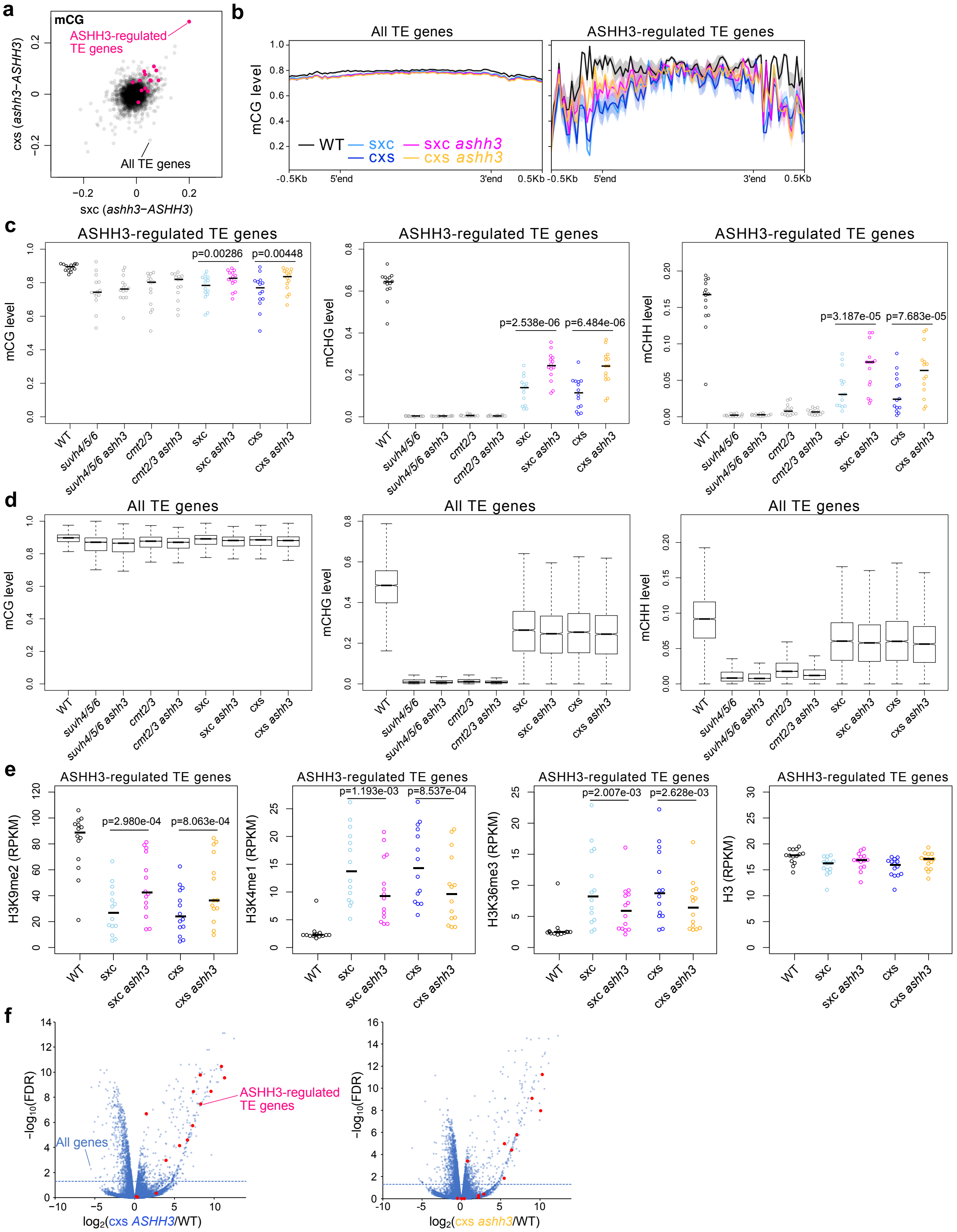

#### Supplementary Fig. 9

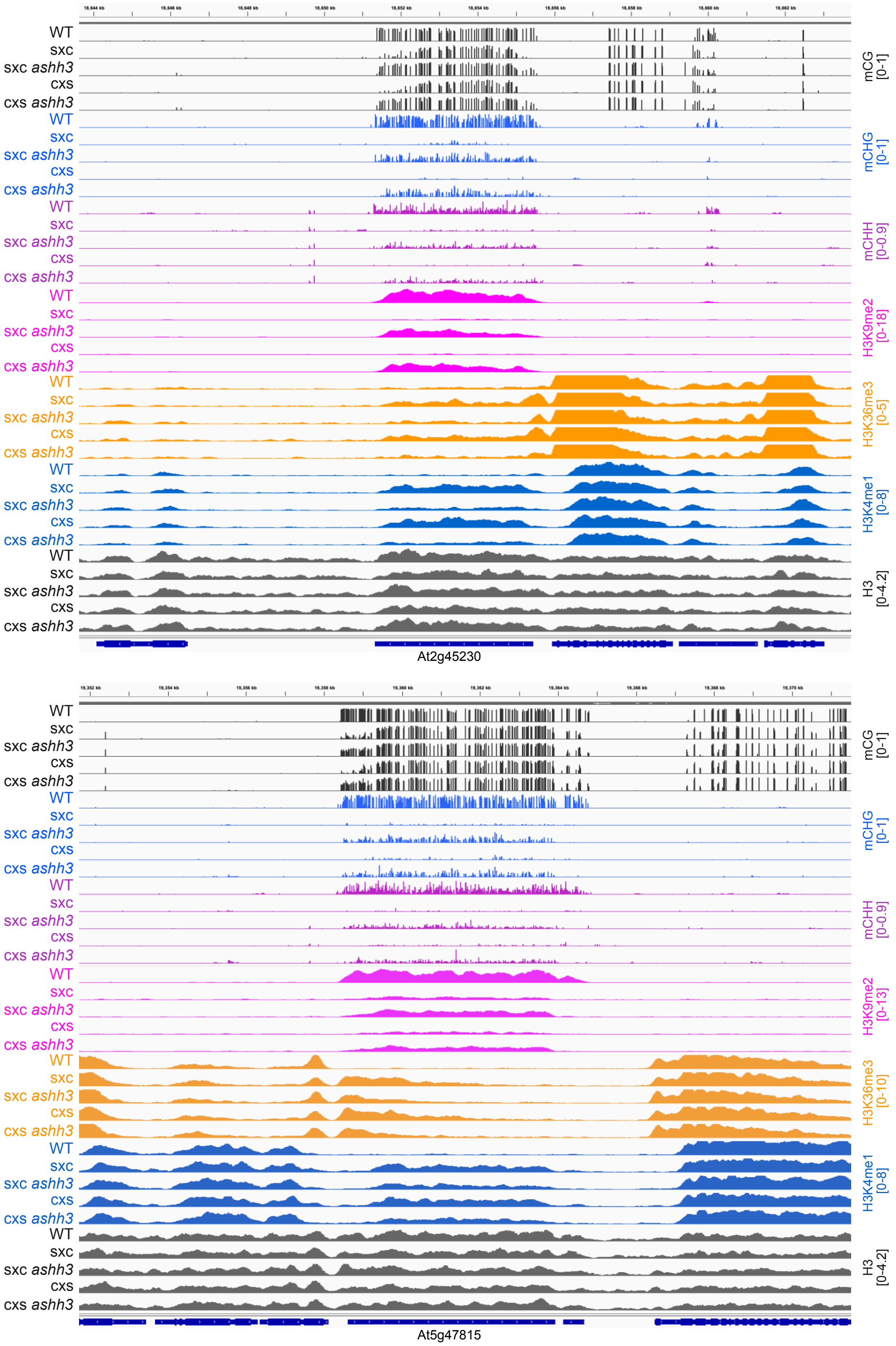
